# A novel menthol-DCMU bleaching method for foraminifera: Generating aposymbiotic hosts for symbiosis research

**DOI:** 10.1101/2024.12.09.627035

**Authors:** Christiane Schmidt, Diana N. Puerto Rueda, Moritz Nusser, Clinton A. Oakley, Xavier Pochon, Marleen Stuhr, Débora S. Raposo, Simon K. Davy

## Abstract

Predicting the response and resilience of coral reefs to climate change can be achieved through better understanding the cellular symbiosis between coral reef holobionts and their associated endosymbiotic algae. Larger benthic foraminifera (LBF) are key calcium carbonate producers, of which two species were investigated for their suitability for menthol bleaching. The LBF *Amphistegina lobifera,* hosting diatoms, and *Sorites orbiculus*, hosting dinoflagellates of the family Symbiodiniaceae. This study aimed to rapidly generate symbiont-free (aposymbiotic) hosts via treatment with menthol and DCMU. The first experiment, Menthol Concentration Comparison (MCC), aimed to find a non-lethal and effective dose for both species. The second experiment, Menthol-bleaching Ecophysiology Assessment (MEA), used a larger sample size of both species to test the response to one concentration 0.19 mmol L^-1^ and measured growth, motility (an indicator for overall fitness) and mortality over a 4-week time frame. Menthol led to an aposymbiotic state in 100% of *A. lobifera* and only minimally impacted its motility and mortality. The method was effective for *S. orbiculus*, where an aposymbiotic state, defined as no visible remains of symbiont cells inside the host at the end of the experimental period, occurred in 66% of specimens of the MCC experiment. Growth was strongly impacted by the bleaching protocol in both species, allowing no new calcite to be formed during the acute exposure. This method can be applied for testing aspects of symbiosis establishment in LBF as well as their potential to take up different symbionts in a short-to medium time frame.

## 1. Introduction

Coral reefs rely heavily on the symbiotic relationship between coral hosts and their endosymbiotic dinoflagellates of the family Symbiodiniaceae (LaJeunesse et al., 2018) to enhance calcification and flourishing in oligotrophic tropical seas (Muscatine and Porter 1977; Morris et al. 2019). Symbiodiniaceae are not only found in corals, but are also associated with other carbonate producers on reefs, such as larger benthic foraminifera (LBF) (Pochon et al. 2004; Fay et al. 2009). Larger benthic foraminifera are important eukaryotic carbonate producers, producing up to 43 Mio. tons of carbonate per year, which represents roughly 0.76 percent of the CaCO_3_ production in the global ocean (Langer et al. 1997). LBF rely on photosynthetic endosymbionts, such as diatoms and dinoflagellates, for their growth and calcification, however the diversity of symbionts reaches beyond those groups to chlorophytes and rhodophytes (Leutenegger 1983; Lee 1998). While they form stable and persistent symbioses with eukaryotic partners, their associations with prokaryotic microbes are flexible and site-specific (Prazeres et al. 2017).

This symbiosis is ecologically significant, yet it has been far less studied compared to the well-documented symbiosis in cnidarians or corals. Hence, our understanding of how foraminifera might respond to anthropogenic impacts, such as climate change, is still limited. This is especially important given that the thermally-induced dysfunction of photo-symbioses on reefs, caused by the climate change-induced increase in frequency and intensity of marine heating events, has far-reaching consequences for ecosystem function and services that are essential for coastal communities worldwide (Hoegh-Guldberg 2004; Wild et al. 2011). Therefore, there is an urgent need to address this knowledge gap in bleaching research involving other calcifiers, and to develop strategies for assisting natural populations to survive climate change. This new field of research is named assisted evolution. In recent years this has been suggested as a strategy for saving coral reefs (van Oppen et al. 2015), for example through the synthesis of new and more thermally-robust host-symbiont pairings via experimental evolution of the symbionts (Scharfenstein et al. 2024). In line with these efforts, our paper provides a methodological approach for rearing aposymbiotic foraminifera, with the ultimate goal of addressing similar research topics to those highlighted for corals and perhaps enhancing the thermal tolerance of foraminifera. This latter strategy could be especially promising given that foraminifera are naturally associated with a wide diversity of symbiotic microalgae.

For many years, a combination of heat or cold stress, dark incubation and/or treatment with the photosynthetic inhibitor 3-(3,4-dichlorophenyl)-1,1-dimethylurea (DCMU), has been used to induce photosymbiont loss, especially in cnidarians (Belda-Baillie et al. 2002; Xiang et al. 2013; Lehnert et al. 2014). However, this approach is time-consuming, often taking several months to create completely aposymbiotic specimens. Therefore, a more rapid and effective method has been developed over the past decade that induces bleaching of corals and sea anemones through a combined incubation with menthol and DCMU (Wang et al., 2012; Matthews et al., 2016; Puntin et al., 2023). Menthol blocks voltage-operated sodium channels in the holobiont, causing local anesthetic effects (Haeseler et al. 2002). The exact biochemical reasons why it induces bleaching in cnidarians are poorly understood, however it causes no or minor discernible impacts on the overall condition of the host organism (Wang et al. 2012; Matthews et al. 2016).

The application of model systems for coral reef research has accelerated in recent years, with, for example, the sea anemone *Exaiptasia diaphana* (commonly referred to as “Aiptasia”) and the upside-down jellyfish *Cassiopea xamachana* helping to elucidate the processes underlying the cnidarian-dinoflagellate symbiosis (e.g. Weis et al. 2008; Medina et al. 2021). Most recently, the menthol bleaching method was applied to a scleractinian coral model, *Galaxea fascicularis*, which is emerging as a particularly amenable organism (Puntin et al. 2023). We currently lack an equivalent model system for coral reef foraminifera, despite their considerable contributions to coral reef productivity and biogeochemical cycling (reviewed in Narayan et al. 2022) and responses to global warming (reviewed in Kawahata et al. 2019). LBF are easy to collect for highly replicated experiments, making them ideal model system candidates. In this study, we tested the potential of two species of common reef-associated LBF, *Amphistegina lobifera* and *Sorites orbiculus*, which host diatoms and dinoflagellates, respectively; the two algal taxa most commonly associated with LBF. Specifically, we aimed to test whether these reef organisms can be bleached using menthol-DCMU incubations without causing increased mortality or hindering growth or motility. By rearing aposymbiotic LBF, we eventually hope to inoculate them with different types of symbionts, to answer questions relating to gshifts in the symbiont community composition in relation to environmental change (‘shuffling’) and host-symbiont specificity.

## 2. Materials and methods

### Sampling and culturing of the foraminifera

The foraminiferan *Amphistegina lobifera* is a tropical to subtropical species abundant on coral reefs worldwide (Langer and Hottinger 2000). Since the opening of the Suez Canal, 150 years ago, Amphisteginids and other LBF have spread throughout the Mediterranean Sea (Zenetos et al. 2008; Weinmann et al. 2013). As the Mediterranean warms, the genus Amphistegina is extending their geographic range westward (Guastella et al. 2019), as their upper thermal tolerance is retained from locations in the Red Sea (Schmidt et al. 2016). For this study, samples were collected from a shallow littoral habitat at Capo Passero, Sicily, Italy, Mediterranean Sea (GPS 36.686667, 15.138278), a location representing, to our knowledge, the ‘invasion front’ (Raposo et al. 2023). A second foraminiferan species, *Sorites orbiculus,* was sourced from the Red Sea, near the Inter-University Institute for Marine Sciences in Eilat, Israel (GPS 29.501866, 34.917488). Samples at each location were collected by snorkelling and removing small-sized seabed rubble at shallow depth (<3 m). Once on shore, seabed rubble was brushed to remove sediment and attached foraminifera. Samples were shipped to New Zealand inside an insulated container containing filtered natural seawater inside Eppendorf vials (volume: 60 ml). Upon arrival, they were maintained in a temperature-controlled room at 23-25°C under white fluorescent light (AQUA-GLO T8 fluorescent bulbs) at 15 µmol photons m^-2^ s^-1^ on a 12:12 h light:dark cycle. The vials in which the LBF cultures were kept were covered with parafilm to limit evaporation and aerated using a small air pump. Specimens were kept in freshly made artificial seawater (Red Sea Salt) at 37-40 psu. For the experiments, they were transferred to a setup filled with local 0.22 µm-filtered seawater sourced from the supply at Victoria University of Wellington, New Zealand.

### 6-week Menthol Concentrations Comparison (MCC) & 33-day High Concentration Trial (HCT)

Preliminary trials aimed to identify a non-lethal dose of menthol to generate aposymbiotic foraminifera. Four different menthol concentrations on the lower spectrum were trialled: 0.05, 0.1, 0.19, and 0.25 mmol l^-1^, using a small sample size (n=3) *per* concentration. This Menthol Concentration Comparison (MCC) experiment was performed on both species using single foraminifera in their jars, monitoring their response with respect to mortality, motility and bleaching, using epifluorescence microscopy, over a 6-week time frame. They were kept seperatly to better observe indivdual responses and track individuals through confocal microscopy. Additionally, a High Concentration Trial (HCT) was performed over 33-days only exposing *A. lobifera* to higher menthol concentrations (0.35, 0.4, 0.45, 0.50, 0.55 and 0.6 mmol l^-1^). This trial was performed to test if any shock responses to the menthol-DCMU treatment occurred at higher concentration, using a small sample size (n=3) *per* concentration, but combining the specimens inside a single jar.

### 4-week Menthol-bleaching Ecophysiology Assessment (MEA) measuring growth and mortality

The menthol-bleaching protocol was adapted from the methods developed by Matthews et al. (2016) for rearing aposymbiotic Aiptasia. The final incubation solution consisted of 0.19 mmol l^-1^ menthol (Sigma Aldrich, NZ) and 5 µmol l^-1^ DCMU ( from a stock solution of 100 mmol l^-1^ dissolved in EtOH; Sigma Aldrich, NZ) in 1 µm-FSW. DCMU functions here as a photosynthetic inhibitor and was added to prevent algal growth and to limit the potential for expelled symbionts to re-enter the host. The foraminifera were incubated in a menthol-DCMU-FSW solution for 6-8 h for five days a week, which was replaced by a FSW-DCMU solution (5 µmol l^-1^ DCMU in 1µm-FSW) for the remaining hours of the day. This 24-h cycle was repeated weekdays for a total of four consecutive weeks, while during weekends samples were kept in the DCMU-FSW treatment solution (5 µmol l^-1^ DCMU in 1 µm-FSW). Epifluorescence microscopy was used to assess bleaching and pseudopodial movement at initial and final time points, and growth was also measured, as described below.

### Experimental design to test menthol bleaching success

The foraminifera were held in screw-capped plastic jars (120 ml) with translucent plastic lids. The jars stood on a plastic grid illuminated at 30 µmol photons m^-2^ s^-1^ on a 12:12 h light cycle under a fluorescent aquarium light source (AQUA-GLO T8 fluorescent bulbs). Temperature control was performed via a water bath set to 25°C (Julabo V8 Thermal pump). Feeding was performed daily in the evening, directly after a water change to the DCMU-FSW solution. A total of 30 µL *Nannochloropsis* feeding concentrate per jar were added, prepared as described in previous culturing experiments (Schmidt et al., 2015).

Each jar contained 50 ml of menthol-DCMU solution in FSW in varying concentrations (see above) or only FSW (controls). In the MCC experiment and the HCT, a single foraminifera was held inside each jar to prevent any stress response from influencing other foraminifera. In the MEA experiments, to allow for statistical evaluation, the number of specimens per jar was increased and varied from 3 to 16, with the smallest sample size used for microscopy and the largest for growth assessments.

### Epifluorescence microscopy

Individual foraminifera from each treatment were examined visually inside standard 6-well cell culture plates filled with fresh 1 µm-FSW. *Amphistegina lobifera* was imaged at x 10 magnification, and *S. oribiulus* at both x 4 (to take an overview picture) and x 20 magnification (close-up). Different magnifications were chosen to best observe the reduction in symbiont density throughout the experiment, using an epifluorescence microscope equipped with an attached camera (Olympus BX63, DP73 Camera). Observations were made at initial and final time points of the MEA experiment and weekly for the MCC experiment. High-resolution fluorescence images (2400 x 1800 Pixel) were acquired using a DAPI filter (excitation 382-393 nm, 417-477 nm emission), which illuminates the symbionts in bright red and the calcium carbonate test in blue. The combination of the epifluorescence lighting and the brightfield setting allowed simultaneous imaging of the symbionts and pseudopodia. During the final image acquisition, additional videos of the pseudopods were filmed, to provide further insight into the physiological effects of menthol treatment and to demonstrate pseudopodial movement as a fitness proxy in completely bleached foraminifera.

### Motility and pseudopodial movement

Motility is used as an indicator of holobiont fitness in foraminifera (Schmidt et al. 2011). It was recorded before each water exchange in all experiments, by documenting the location of each *A. lobifera* specimen on the wall (vertical) or on the bottom of the culturing jar (horizontal). The data for *A. lobifera* were used to calculate the percentage motility *per* jar and *per* treatment. Cultures were examined daily on weekdays in both experiments. Motility of *S. orbiculus* in the same experiments was not measured.

### Growth and survivorship

For growth measurements, the initial and final size of each specimen were imaged using a compact stereo microscope (ZEISS Stemi 305 with Zen software). From these two measurements, the maximum diameter of each foraminifera was measured (ImageJ software). To obtain growth rates *per* individual, specimens *per* jar were ordered according to their size (from smallest to largest individual) and data were ordered to calculate individual growth rates *via* the formula in Schmidt et al. (2011).

Mortality of benthic living foraminifera is typically assessed by recording vital cytoplasm colour (Bernhard 2000). In this study, however, fully bleached foraminifera often lacked cytoplasm colour and we employed a combination of methods to assess mortality, such as lack of motility, cytoplasm colour and extended pseudopodial nets. Mortality was only systematically assessed using the paraments above in the MEA 4-week experiment, in the MCC 6-week expeirment and the HTC treatment, motitly was used to investigate the effect of the bleaching treatment on fittness of the holobiont. After the MEA 4-week experiment a few individuals of *S. orbiculus* were overgrown by algae at the which were regarded dead in this case. This may have led to a slight increase in their mortality rate of this species compared to *A. lobifera*.

### Statistics

Statistical analyses and graphs were performed in RStudio, using R Version 4.3.0 and the packages tidyverse and ggprism (R Core Team 2023). For growth data in the MEA experiment, the non-parametric Mann-Whitney U test was used to test for differences between treatments (menthol *vs*. control), as the data were not normally distributed nor met the assumption of homogeneity of variances. To analyze the motility data in the MEA 4-week experiment, we used a generalized linear model (GLM) for binomially distributed data with a logit link function, in combination with a generalized estimating equation (GEE) approach to account for the temporal correlation between repeated measurements on the same individuals. Specifically, we utilized the geeglm function within the geepack R package to implement the GEE model (Højsgaard et al., 2006). To address temporal correlation, we incorporated an autoregressive correlation structure of the first order (AR-1) into the model. The predictor variables in our model comprised “treatment” and “day of experiment”, including the interaction term. To assess differences between model parameters, we employed the Wald statistic. The motility of the 6-week MCC (Menthol Concentration Comparision) experiment and the HCT (High Concentration Trial) was not statistically evaluated as sample size were small (n=3).

## 4. Results & Discussion

### 6-week Menthol Concentration Comparison (MCC) & 33-day High Concentration Trial (HCT)

The visual comparison of epifluorescence microscopy images between the different menthol concentrations used for *Amphistegina lobifera* (Figure 1) and *Sorites orbiculus* (Figure 2), showed that menthol bleaching was effective on LBF. For *A. lobifera*, the visually strongest reduction occurred between day 16 to day 33 (Figure 1), and bleaching occurred also at the lower menthol concentrations. For *S. orbiculus*, there were individual differences between the specimens (see specimen #2 was not fully aposymbiotic, Figure 2, conc. 0.25 mmol l^-1^). However, most specimens visually lost their symbionts within the 6 week timeframe (Figure 1,2). Motility, the climbing of *A. lobifera* on the vertical walls of the jar from a flat surface was additionally assessed to observe the holobionts respons to the menthol stressors. The motility data showed no clear trend (Figure 3) in any of the treatments, hence the third highest dose of 0.19 mmol l^-1^ was considered safe for the foraminifera (Figure 3).This concentration has been used successfully with the cnidarian model Aiptasia (Matthews et al. 2016). Therefore, we decided to apply the dose 0.19 mmol l^-1^ ,for the 4-week MEA experiment to access the influence of the menthol also on growth rate in a larger sample size.

**Figure 1.**
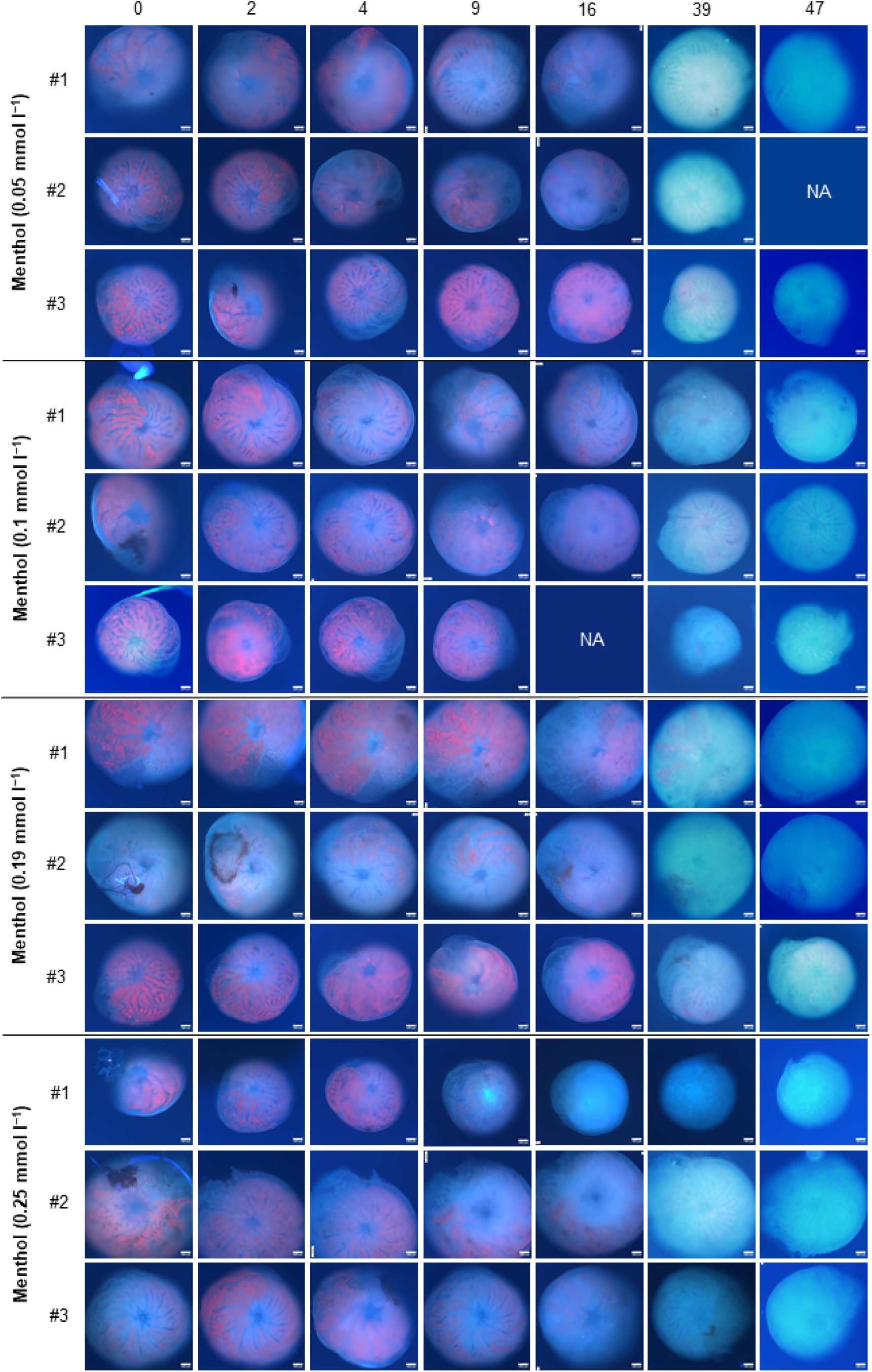
Epifluorescence microscopy images of all *Amphistegina lobifera* individuals during the 6-week MCC (Menthol Concentration Comparision) experiment at seven different time points and different menthol concentrations. Different menthol concentrations are depicted by data of experiment 0.05, 0.1, 0.19, 0.25 mmol l-1, Scale bars are 100 µm.

**Figure 2.**
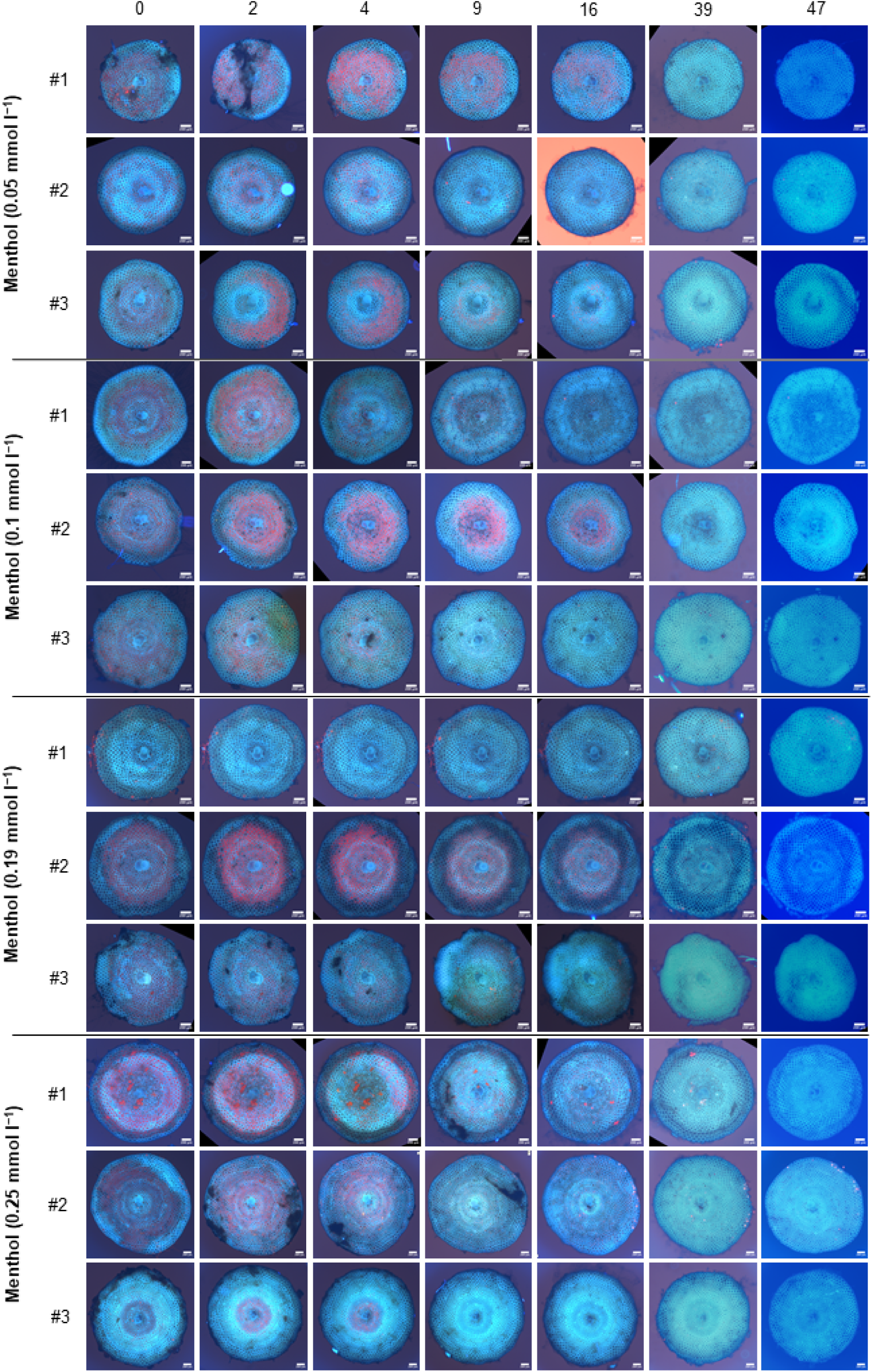
Epifluorescence microscopy images of all *Sorites orbiculus* individuals during the 6-week MCC (Menthol Concentration Comparision) experiment at seven different time points and for different menthol concentrations. Different menthol concentrations are depicted by data of experiment 0.05, 0.1, 0.19, 0.25 mmol l-1, Scale bars are 200 µm.

**Figure 3.**
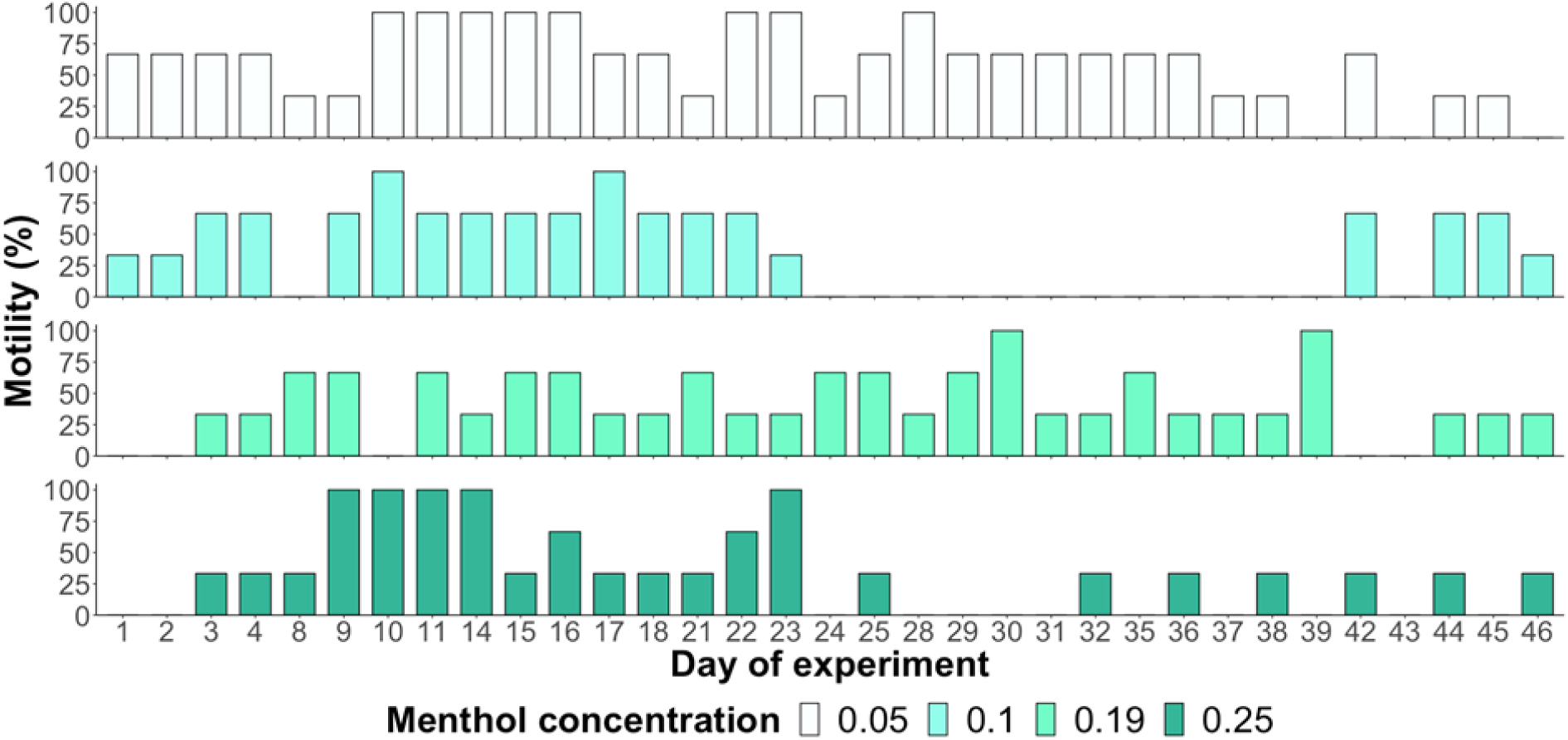
Motility of *Amphistegina lobifera* during the 6-week MCC (Menthol Concentration Comparision) experiment, calculated as the number of individuals on the wall of the jar divided by the total number in the jar. n = 3 *per* concentration. Menthol concentration given in mmol l^-1^.

The HCT High Concentration Trial (up to 0.6 mmol l^-1^) was performed on *A. lobifera* because it has also been tested previously on *Aiptasia*, though it caused 50% mortality in this anemone. During the experiment we did not systematically investigate mortality, but we investigated the fitness of the holobiont through assessment of motility (Figure 4). At the end of the 33-day trial period motiltiy was with <25-50% rather low, and foraminifera seemed more stressed than in the 0.19 mmol l^-1^dose, which was then regarded a save dose to perfom a larger experiment with minimal mortality and still yield effective results.

**Figure 4.**
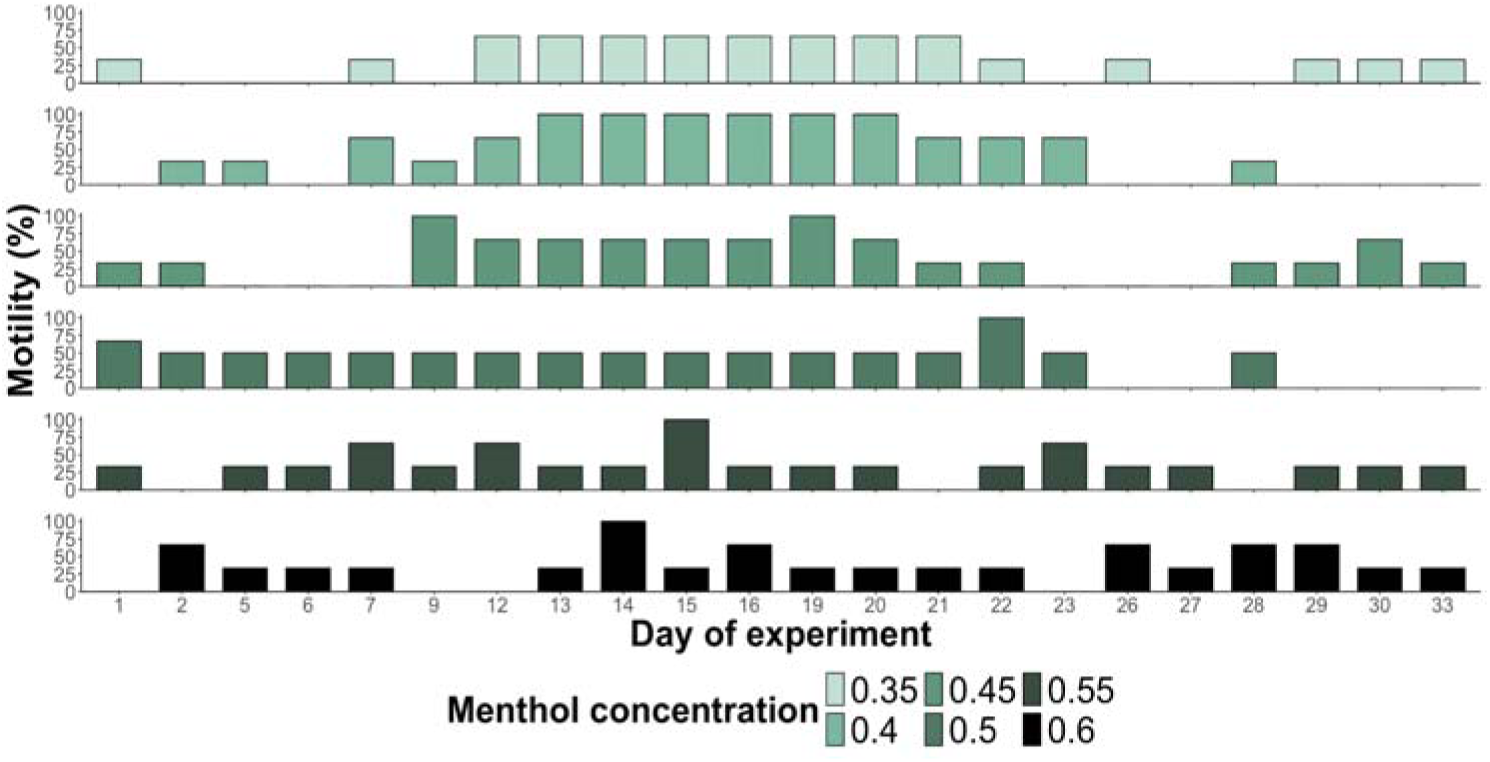
Motility of *Amphistegina lobifera* during the HCT (High Concentration Trial) over 33-days. Motility is calculated as the number of individuals on the wall of the jar divided by the total number in the jar. n = 3 *per* concentration. Menthol concentration given in mmol l^-1^.

### 4-week Bleaching Ecophysiology Assessment (MEA) measuring growth and mortality

The repeated menthol treatment at a concentration of 0.19 mmol l^-1^ induced symbiont loss in the MEA experiment; in many specimen, no symbionts where visibly remaining using confocal microscopy at the end of the experiment (Figure 5A). For *A. lobifera*, all specimens that were individually assessed microscopically lost 100% of their symbionts (n = 3) at the end of the experiment. For *S. orbiculus*, two out of three specimen (66.66%) became complelty aposymbiotic. The remaining one third (33.33%) still hosted autofluorescent symbiont cells upon final observation, as already observed in the MCC experiment in a few cases. The number of cells remaining, however, was extremely small when compared to cell counts of Symbiodiniaceae in the LBF *Marginopora vertebralis*, which are typically in the order of 3.5-5 x 10^4^ cells per mm^2^ surface area (Reymond et al. 2013). This led us to conclude that substantial symbiont loss had occurred, which would be essential for further experimental work. Hence, we suggest individual selection and checking of putatively aposymbiotic specimens in the confocal microsopy is needed before subsequent re-inoculation experiments to be working with 100% aposymbiotic inviduals.

**Figure 5.**
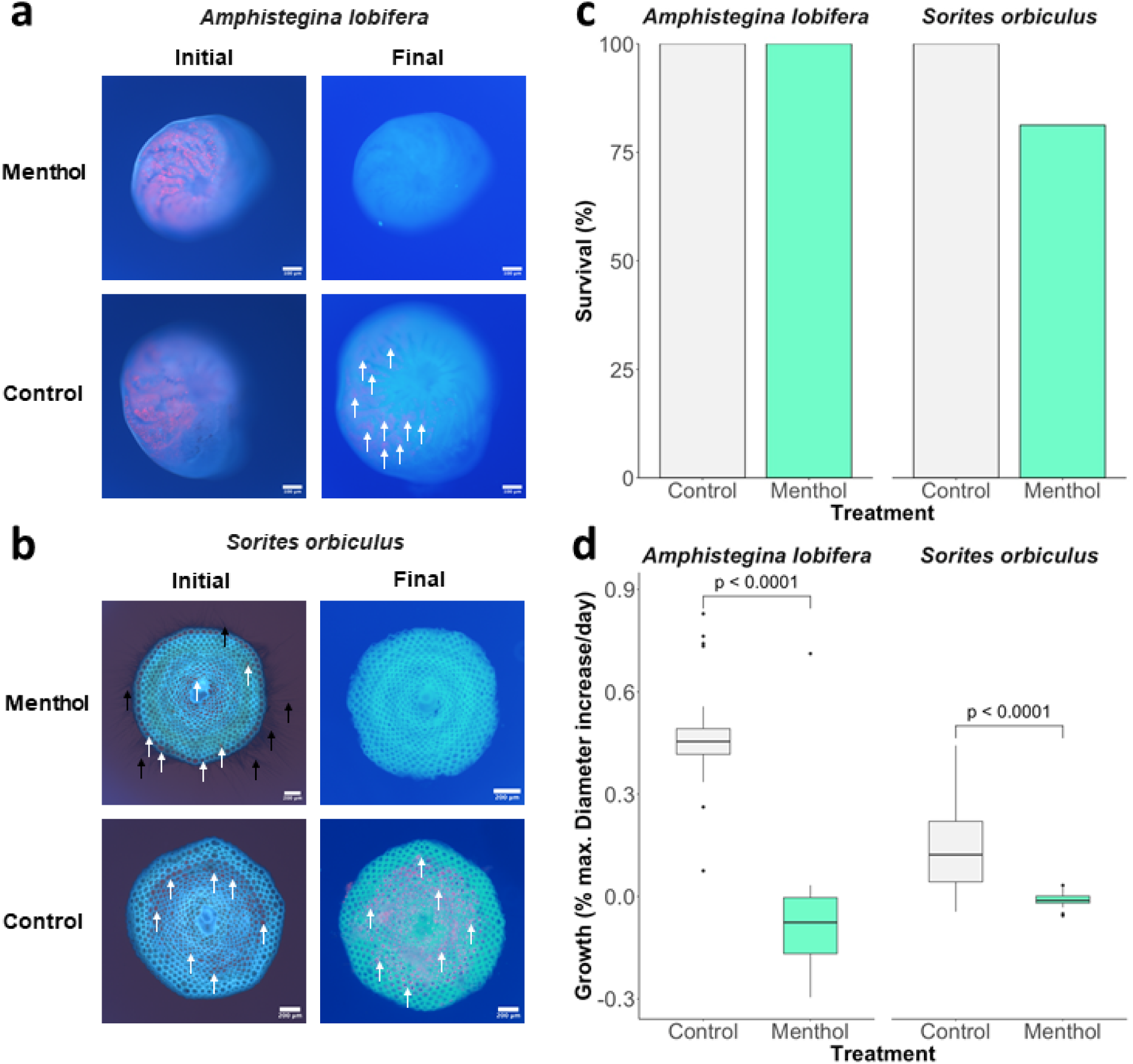
Effects of menthol-DCMU exposure after the MEA (Menthol-bleaching Ecophysiology Assessment) 4-week experiment on the foraminiferan species *Amphistegina lobifera* and *Sorites orbiculus*. **(a-b)** Epifluorescence microscopy images of initial and final assessments showing symbiont loss (selected from n=3 per treatment). Scale bars for *Amphistegina lobifera* and *Sorites orbiculus* are 100 µm and 200 µm, respectively. Fluorescing symbiont cells in the host cell are marked with white arrows, while pseudopodia of the foraminifera are indicated by black arrows. **(c)** Survivorship after the MEA 4-week experiment **(d)** Outlier box plots of growth data given as % diameter increase *per* day. Significant differences between control and menthol treatment of the species is indicated by p values (Mann-Whitney U test, for *A. lobifera* n = 63 for *S. orbiculus* n = 32).

Survivorship remained high (Figure 5B) in both species at the applied concentration, which was tested before too low to cause lethal effects. However, both species were stressed during the menthol bleaching process as reflected in their neutral growth rates. Growth rates were significantly different in both species between the control and menthol treatment (Mann-Whitney U test, Table 1, Figure 5C), indicating that menthol can lower growth rates during the repeated cycle of applying menthol for bleaching in both species. While growth was in the normal range for the controls (Figure 5C), no growth occured in the menthol-DCMU bleaching treatment in either species, clearly indicating that they were under physiological stress. We speculate that the lack of symbiont-derived photosynthate was probably limiting calcification, which has been observed in reef-scale assessments that link coral bleaching with reduced coral growth (Bove et al. 2020; Davis et al. 2021). In this experimental setting there is another possible explaination, that negative growth rates of *A. lobifera* were likely caused by shell breakage, in particular in the menthol-DCMU treatment, as the control treatment showed growth rates in the normal range, which is comparable to previous studies (Schmidt et al. 2016). For *S. orbiculu*s, growth was generally low (Figure 5C), despite 25°C being its optimal culturing temperature for calcification (Kinoshita et al. 2021). This could have to do with that both species had to be kept under the same light intensitiy, which could likely be stimulated in *S. orbiculus* when light levels are a bit higher more optimal for *Symbiodinium* symbionts.

**Table 1.** Statistical results of the Mann-Whitney U Test on growth rates comparison after the MEA (Menthol-bleaching Ecophysiology Assessment) 4-week experiment.

| Species | Compared groups | n (control) | n (menthol) | U | p-value |
| --- | --- | --- | --- | --- | --- |
| *Amphistegina lobifera* | Growth rate ~ Treatment | 32 | 31 | 436 | <0.001 |
| *Sorites orbiculus* | Growth rate ~ Treatment | 16 | 16 | 102 | <0.001 |

Pseudopods, also known as reticulopodia, are key structures in foraminifera, enabling them to move, attach to surfaces, and capture food (Travis et al. 2002). Throughout our microscopy observations, pseudopods were occasionally visible in both the menthol and the control treatments. Clearly visible pseudopods were imaged (Figure 6A-C) and videos of foraminifera in the aposymbiotic state with pseudopods are available as part of the data set (deposited at https://www.pangaea.de/). Due to the intermittent nature and variability of pseudopods in foraminifera (Greco et al. 2023), a better indicator of holobiont fitness is motility. This parameter could be assessed in *A. lobifera* as this species is generally very active in comparison to other benthic foraminifera, such as *S. orbiculus* (C.S. unpublished observations) and *P. calcariformata* (Schmidt et al. 2015). Motility is a direct indicator of active pseudopodial movement and does not require detailed microscopic observations of each individual and is known to decrease in response to thermal stress (Schmidt et al. 2011; Stuhr et al. 2018).

**Figure 6.**
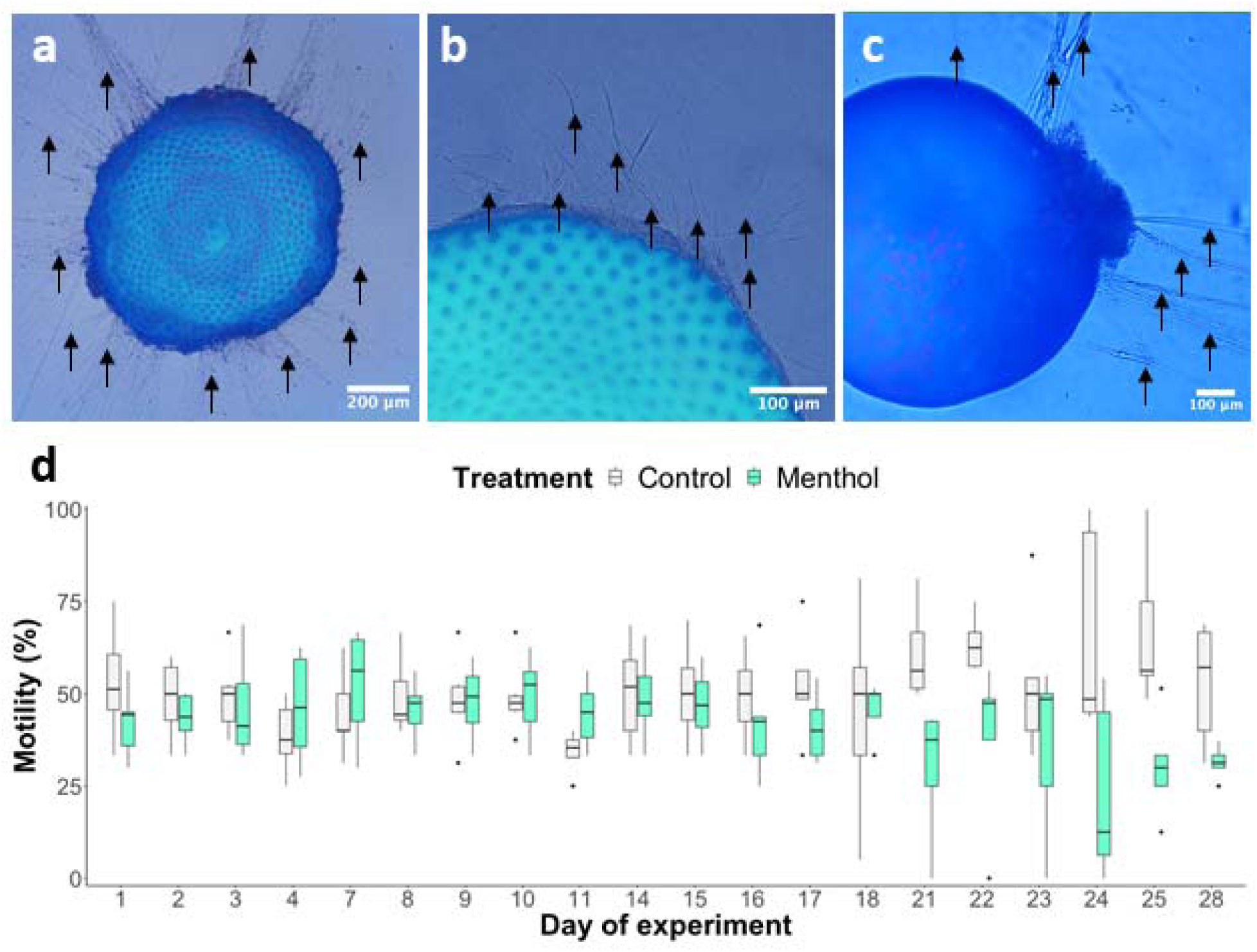
Pseudopodia of selected menthol-bleached and control specimens of *Amphistegina lobifera* and *Sorites orbiculus*, and motility of *A. lobifera* during the MEA (Menthol-bleaching Ecophysiology Assessment) 4-week experiment (a) Extended pseudopodial net of *S. orbiculus* in the control treatment (b) after the menthol-DCMU treatment (c) *A. lobifera* from the control treatment. Black arrows indicate pseudopodia. (d) Outlier boxplot showing changes of motility in *A. lobifera* over time (n = 155 *per* time point, grey boxes = controls and turquoise boxes = menthol treatment).

Motility was not impaired at concentration of 0.19 mmol l^-1^ menthol (Fig. 6D), however after day 21, motility in the control treatment was higher than in the menthol treatment. On the final day of the experiment (day 28), motility was significantly lower in the menthol treatment than the control treatment (Wald statistics, Table 2, p < 0.001). Furthermore, there was a significant interaction between treatment and day-of-experiment (Wald statistics, Table 2, p < 0.001). As expected due to the anaesthetic properties of menthol (Galeotti et al. 2001; Haeseler et al. 2002), repeated treatment led to reduced motility. We assume that menthol had a relaxing effect on pseudopodial activity. Relaxation of tentacles and unresponsiveness due to menthol has been observed in a range of different marine species, such as pteropods (Yamazaki et al. 2024) and anemones (Matthews et al. 2016).

**Table 2.** Statistical results of the GLM (General Linear Model) for the motility data of the MEA (Menthol-bleaching Ecophysiology Assessment) 4-week experiment including the Wald Statistic.

| Predictor variable | df | Wald Chi-Square | p-value |
| --- | --- | --- | --- |
| Day of experiment | 19 | 15 | 0.73 |
| Treatment | 1 | 19.7 | <0.001 |
| Treatment vs. Day of experiment | 19 | 58.7 | <0.001 |

## 5. Conclusion

Our findings demonstrate that large benthic foraminifera (LBF) hosting diatoms and dinoflagellates can be experimentally induced to lose their symbionts (aposymbiosis) using a combination of menthol-DCMU as bleaching agents. This process is achievable within 4–6 weeks without inducing lethal stress to the holobionts. Yet, we observed effects on their fitness as demonstrated by a lack of growth and reduced motility compared to the controls. It is now important to test whether we can re-inoculate these menthol-treated LBF with different symbionts, and their ecophysiological health (e.g. normal growth & calcification etc.) can be reestblished when they have taken up symbionts again. The insights gained from the experimental application of aposymbiotic foraminifera have broad implications for the field of coral reef ecology, including our understanding of adaptive strategies such as the potential acquisition of novel, more thermally tolerant symbionts that could benefit these important organisms in the face of climate change.

## 6. Competing interests

The authors declare no competing or financial interests.

## Acknowledgements

Funding to conceive the study and conduct the research stay at University of Wellington was given to C.S. through a DAAD (German Academic Exchange Service) short-term postdoctoral grant. The data analysis and personal costs was supported by the DFG (German Research Foundation) grant SYMBIO-AID which was an Individual Research Grant given to C.S. (SCHM 3395/3-1). A grant from the Marsden Fund of the Royal Society Te Apārangi (No. 19-VUW-086) was awarded to S.K.D. and C.A.O, who provided funding for consumables and lab support at Victoria University of Wellington. During sample collection for this study, M.S. was supported by the Minerva Foundation and D.S.R. by the BMBF (Grant Number 03F0820A).

## Summary statement

Novel application of a menthol-based method for producing symbiont-free benthic foraminifera for further study of symbiosis establishment and function in these ecologically important symbiotic organisms.

## 7. Author contributions

M.S. and D.S.R provided samples, C.S., X.P., C.A.O., and S.K.D designed and conceived the experiments. C.S., M.N. and D.N.P. R carried out the experiments and subsequent data analysis. C.S., M.S., D.S.R., X.P., C.A.O., D.N.P.R. and S.K.D. wrote and edited the manuscript.

## 8. Data availability

Datasets are uploaded to the PANGAEA (deposited at https://www.pangaea.de/).

## Notes

https://www.pangaea.de/

### Competing Interest Statement

The authors have declared no competing interest.

